# Features of smaller ribosomes in Candidate Phyla Radiation (CPR) bacteria revealed with a molecular evolutionary analysis

**DOI:** 10.1101/2022.01.07.475337

**Authors:** Megumi Tsurumaki, Motofumi Saito, Masaru Tomita, Akio Kanai

**Author notes:** Corresponding Author Akio Kanai, PhD Institute for Advanced Biosciences, Keio University, Tsuruoka, Yamagata 997-0017, Japan.

## Abstract

The Candidate Phyla Radiation (CPR) is a large bacterial group consisting mainly of uncultured lineages. They have small cells and small genomes, and often lack ribosomal proteins L1, L9, and/or L30, which are basically ubiquitous in ordinary (non-CPR) bacteria. Here, we comprehensively analyzed the genomic information of CPR bacteria and identified their unique properties. In the distribution of protein lengths in CPR bacteria, the peak was at around 100–150 amino acids, whereas the position of the peak varies in the range of 100–300 amino acids in free-living non-CPR bacteria, and at around 100–200 amino acids in most symbiotic non-CPR bacteria. These results show that CPR bacteria have smaller proteins on average, like symbiotic non-CPR bacteria. We found that ribosomal proteins L28, L29, L32, and L33 are also deleted in CPR bacteria, in a lineage-specific manner. Moreover, the sequences of approximately half of all ribosomal proteins in CPR differ, in part, from those of non-CPR bacteria, with missing regions or specific added region. We also found that several regions of the 16S, 23S, and 5S rRNAs are lacking in CPR bacteria and that the total predicted length of the three rRNAs in CPR bacteria is smaller than that in non-CPR bacteria. The regions missing in the CPR ribosomal proteins and rRNAs are located near the surface of the ribosome, and some are close to one another. These observations suggest that ribosomes are smaller in CPR bacteria than in free-living non-CPR bacteria, with simplified surface structures.

## INTRODUCTION

Studies of microbial communities based on sequence analyses of DNA extracted directly from an environment without culturing the microorganisms, such as metagenomic analyses or 16S rRNA gene sequencing, have revealed large numbers of microbial lineages that do not belong to known classification groups (Castelle and Banfield 2018). The Candidate Phyla Radiation (CPR) is a monophyletic group in the bacterial domain that is mainly composed of uncultured lineages (Brown et al. 2015; Hug et al. 2016) (Supplementary Figure S1). At least 74 candidate phyla belonging to the CPR bacteria have been reported, and these bacteria are widely distributed in various environments, including soil, sediments, groundwater, fresh water or the human oral cavity (Wrighton et al. 2012; Kantor et al. 2013; Rinke et al. 2013; Brown et al. 2015; Luef et al. 2015; Anantharaman et al. 2016). None of these bacteria have been cultured, except Saccharibacteria derived from the human oral cavity. When prokaryotic genomes are clustered according to the presence or absence of 921 widely distributed protein families, CPR bacteria are clearly seen to have evolved separately from other bacteria (Meheust et al. 2019). On a phylogenetic tree of the three domains of life (Bacteria, Archaea, and Eukaryota) constructed by Hug et al. based on ribosomal proteins, CPR diverged at the base of the bacterial domain, forming a clade that is separated from all other bacteria (Hug et al. 2016). However, the reliability of the branches at deep positions is considered insufficient on that phylogenetic tree. In contrast, in a bacterial phylogenetic analysis by Coleman et al., CPR is a sister group of the phylum Chloroflexi in the Terrabacteria group, which consists of multiple phyla (Coleman et al. 2021). Therefore, no conclusions have been reached regarding the phylogenetic position of CPR bacteria. The scale of the diversity of CPR bacteria is also unclear. The phylogenetic analysis of Hug et al. (2016) suggested that CPR bacteria account for about half of all bacterial diversity. In contrast, Parks et al. (2017) predicted that CPR bacteria account for 26.3% of all bacterial diversity. In either case, the presence of CPR cannot be ignored in any discussion of bacterial diversity and evolution.

Despite the magnitude of their phylogenetic diversity, CPR bacteria have some common characteristics. First, many CPR bacteria are small cells, reflected in the fact that they are abundant in samples that have passed through filters with a pore size of 0.2 µm (Miyoshi et al. 2005; Brown et al. 2015; Luef et al. 2015) and in electron microscopic observations (He et al. 2015; Luef et al. 2015). Their genomes are also small, and most genomic sequences obtained with metagenomics or single-cell genomics indicate sizes of ≤1.5 Mb, which is close to those of symbiotic bacteria. These genomes lack genes of important metabolic pathways, such as the tricarboxylic acid cycle and amino acid and nucleotide biosynthesis pathways (Kantor et al. 2013; Nelson and Stegen 2015; Danczak et al. 2017; Suzuki et al. 2017; Castelle et al. 2018). The characteristics of CPR bacteria, such as their small genomes and lack of biosynthetic capacity, are similar to those of symbiotic organisms. The lifestyles of many CPR bacteria have not been reported, except for Saccharibacteria, which attaches to a bacterium of the genus *Actinomyces* (He et al. 2015), and a lineage found to live in the cytoplasm of *Paramecium bursaria* (Gong et al. 2014).5

The distinctiveness of CPR bacteria also extends to their ribosome-related genes. The bacterial ribosome is composed of the 30S small subunit (SSU) and the 50S large subunit (LSU), which consist of multiple ribosomal proteins and three rRNAs (16S in the SSU, and 23S and 5S in the LSU) (Schuwirth et al. 2005). In *Escherichia coli*, the SSU contains 21 proteins and the LSU contains 33 proteins (Kaczanowska and Ryden-Aulin 2007). Some CPR bacteria contain one or more introns in their 16S rRNA and 23S rRNA genes, although these are very rare in other bacteria (Brown et al. 2015). All CPR bacteria lack the ribosomal protein L30. L1 is absent in some species of Parcubacteria (a CPR subgroup); L9 is absent in most Microgenomates (a CPR subgroup) and other candidate phyla Dojkabacteria, WWE3, and Saccharibacteria (Brown et al. 2015). Although these ribosomal proteins are not considered essential for bacterial survival, they are known to occur widely in bacteria other than some symbionts (Yutin et al. 2012; Nikolaeva et al. 2021). Because ribosomes have fundamental functions associated with basis life processes, their structures are considered highly conserved. However, in recent years, large-scale studies of the distribution of ribosomal proteins have shown that bacteria with small genomes often lack certain ribosomal proteins. It has been proposed that the structures near the surface of the ribosome vary in regions that were acquired in the “late phase” of the molecular evolution of the ribosome (Yutin et al. 2012; Nikolaeva et al. 2021). Generally, reduced genomes are found at the level of a single phylum or genus, whereas in CPR bacteria, small genomes occur throughout a large clade containing multiple phyla (Brown et al. 2015; Castelle et al. 2018). Therefore, understanding the structure of the CPR ribosome will not only clarify its origin, but is also important in any discussion of the diversity and evolution of bacterial ribosomes in general. However, in characterizing the ribosomes of CPR bacteria, although missing proteins have been identified over a wide range of lineages, little is known about the sizes and structures of individual proteins.

In recent years, the number of prokaryote genomes registered in public databases has steadily increased. CPR bacteria are no exception, and in addition to many partial and draft genomes, approximately 70 complete genomes have been registered. In this study, we attempted to characterize the ribosomes of CPR bacteria based on a comparison of 69 complete and 828 draft genomes of CPR bacteria with known non-CPR bacterial genomes. The size distribution of all bacterial proteins predicted from each individual genome shows that CPR bacteria, like some parasites, have smaller proteins, on average. Moreover, some CPR bacteria lack several ribosomal proteins that have not been previously noted. A comparison of the amino acid sequences of ribosomal proteins revealed regions that are absent only in CPR bacteria and regions that are present only in CPR bacteria. A comparison of the ribonucleotide sequences of each rRNA also revealed RNA regions that are only absent in CPR bacteria or only present in CPR bacteria. In the three-dimensional ribosomal structure, the missing regions in the rRNAs and ribosomal proteins are unevenly located near the ribosome surface. These results suggest that the ribosomes of CPR bacteria are small, with simplified surface structures.

## RESULTS AND DISCUSSION

### Smaller protein sizes in CPR bacteria

To clarify the characteristics of CPR bacterial genomes using as many examples as possible, we first comprehensively compared the gene lengths of CPR bacteria with those of other well-known (non-CPR) bacteria. For this purpose, 69 complete and 828 draft genomes of CPR bacteria and 1,661 complete genomes of non-CPR bacteria were collected. The non-CPR bacteria were divided into 167 parasitic or symbiotic lineages (non-CPR symbiotic) and 1,494 other lineages (non-CPR free-living) (Supplementary Tables S1–S4). Because symbiotic bacteria often have small genomes or reduced biosynthetic capacities, in common with CPR bacteria (Castelle et al. 2018), comparing CPR bacterial genomes with symbiotic bacterial genomes can highlight the similarities and differences among reduced genomes. Based on 43 single-copy genes, the genomes of CPR bacteria were estimated to range from approximately 0.3 to 1.7 Mb. In contrast, the genome sizes of free-living and symbiotic non-CPR bacteria were calculated to range in size from approximately 1.3 to 13.0 Mb and from 0.3 to 8.8 Mb, respectively. This confirmed that CPR bacterial genomes are as small as those of symbiotic bacteria, as reported previously (Castelle et al. 2018).

A total of 721,344 proteins (191–1,723 proteins per genome) of CPR were identified with the gene-finding program Prodigal (Hyatt et al. 2010), and 5,388,587 proteins (237–10,234 proteins per genome) of the non-CPR bacteria were identified according to the genome annotations in National Center for Biotechnology Information (NCBI) RefSeq. Figure 1A and 1B shows the distribution of protein lengths in the 69 complete genomes of CPR bacteria and in representative genomes of non-CPR bacteria (70 free-living and 70 symbiotic species), respectively, along with density curves (see Supplementary Figure S2 for the distribution of protein lengths in all the genomes used in this study). In the distribution of protein lengths of CPR bacteria, the peak was at around 100–150 amino acids (aa), whereas the position of the peak varied in the range of 100–300 aa in free-living non-CPR bacteria (Figure 1A). Among the free-living non-CPR bacteria, some genomes showed multimodal distributions of protein length, with one peak at approximately 150 aa and another at approximately 300 aa (e.g., *Bacteroides fragilis* and *Bacillus subtili*s). Compared with the distribution of protein lengths of free-living non-CPR bacteria, CPR bacteria tended to have a higher proportion of proteins of ≤250 aa. These results show that CPR bacteria not only have smaller genomes, but also smaller proteins, on average. Among the non-CPRs, the peak in the protein length distribution was around 100–200 aa in most symbiotic bacteria, which is closer to that of CPR bacteria than to that of other (free-living) non-CPR bacteria (Figure 1B). For example, the distributions of TC1 (an endosymbiont of *Trimyema compressum*), *Spiroplasma mirum, Coxiella burnetii*, and *Wolbachia* sp., which are intracellular endosymbionts or pathogens with genomes of <1.6 Mb, were highly skewed to the left, with sharp peaks at around 100 aa.

**Figure 1.**
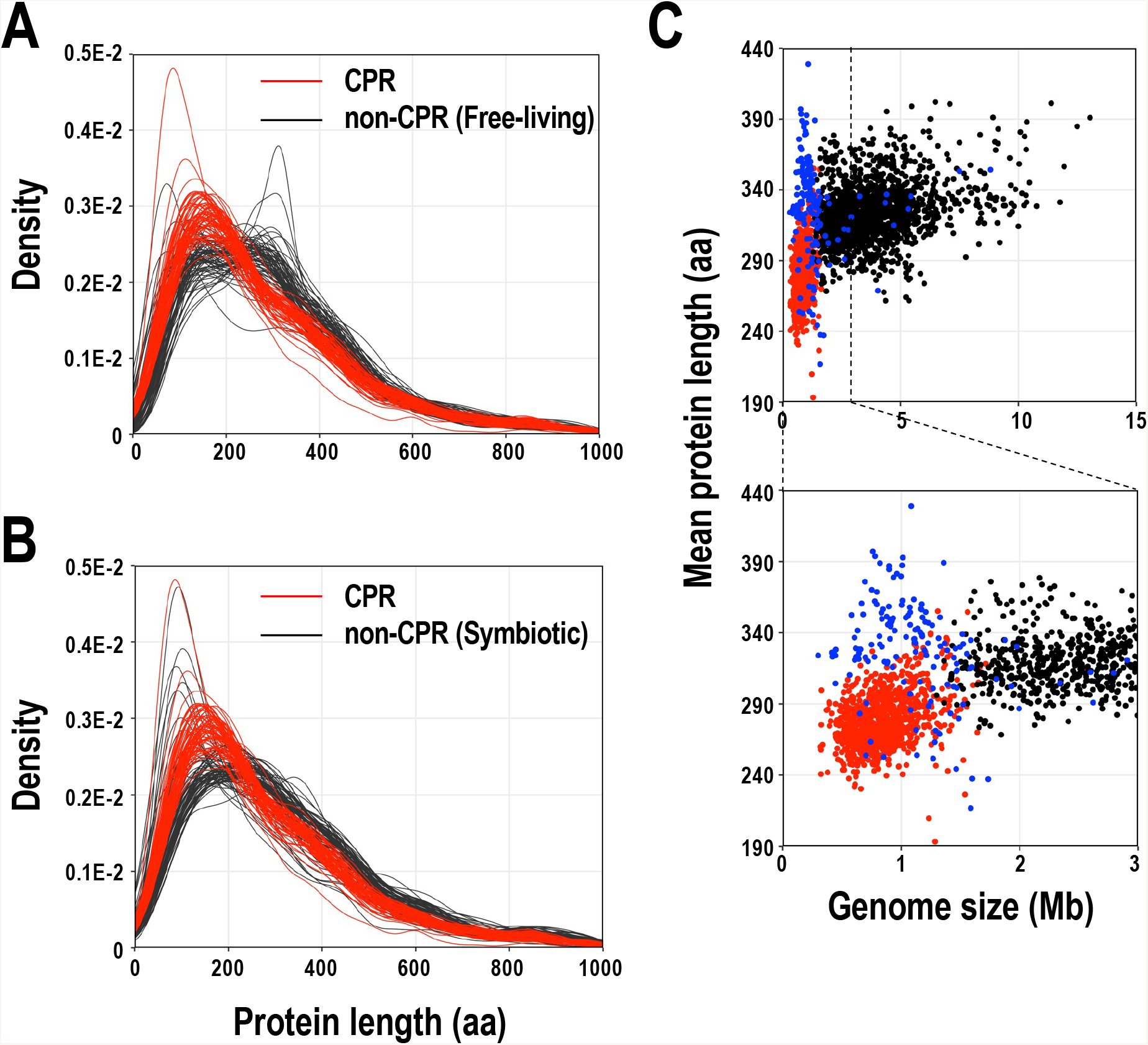
Comparison of lengths of proteins encoded in complete CPR and non-CPR genomes. (A) Comparison of the lengths of proteins from CPR and non-CPR (free-living species) bacteria. (B) Comparison of the lengths of proteins from CPR and non-CPR (symbiotic species) bacteria. Distributions of lengths of proteins are shown as a density curve using the deduced amino acid sequences of the protein genes in 69 CPR bacteria (red line) and 140 representative non-CPR bacteria (70 free-living lineages and 70 symbiotic lineages; each black line). (C) CPR bacteria have smaller genomes with shorter proteins. We used 897 CPR bacterial genomes (69 complete and 828 draft genomes; red dots), 167 symbiotic non-CPR bacterial genomes (blue dots), and 1,494 free-living non-CPR bacterial genomes (black dots) in the analysis. The lower panel is an enlarged view of part of the upper panel.

Plots of the mean protein length per genome (Figure 1C) showed similar results. The mean protein length per CPR bacterial genome (193–355 aa, mean 279 aa) was smaller than that of free-living non-CPR bacteria (262–402 aa, mean 325 aa; *p* < 0.01, Mann–Whitney *U* test). Although most symbiotic non-CPR bacteria have small genomes like CPR, the mean protein length in their genomes (217–429 aa, mean 329 aa) was larger than that of CPR (*p* < 0.01, Mann–Whitney *U* test). As an exception, some non-CPR bacteria, such as TC1 (217 aa on average), *Coxiella* sp. (237 aa long on average), and *Spiroplasma citri* (237 aa long on average), had particularly small mean protein lengths. Interestingly, the mean protein length correlated weakly with genome size in the CPR bacteria (R = 0.36, *p* < 0.01) and free-living non-CPR bacteria (R = 0.30, *p* < 0.01), but no correlation was observed in symbiotic non-CPR bacteria.

### Lack of certain ribosomal proteins in the CPR bacterial lineages

The CPR genomes examined in this study were classified into 65 candidate phyla with reference to the NCBI taxonomy (Schoch et al. 2020) and a previously reported phylogenetic tree (Hug et al. 2016). Fifty-three of these phyla belonged to two previously proposed superphyla: Parcubacteria (OD1; 39 phyla) and Microgenomates (OP11; 14 phyla). The other 12 phyla were tentatively classified into two subgroups based on the phylogeny. The phyla closely related to Parcubacteria were classified as “phylum group I”, and the phyla closely related to Microgenomates as “phylum group II” (Supplementary Figure S1 and Supplementary Table S1A). To comprehensively extract the 54 ribosomal proteins widely distributed in bacteria from the genomes of CPR bacteria, we obtained known ribosomal protein sequences registered in the NCBI Clusters of Orthologous Groups of proteins (COG) database (Galperin et al. 2021b) (Supplementary Table S5) and domain data for the ribosomal proteins registered in the Pfam database (Supplementary Table S6; (Mistry et al. 2021). Using these data, the ribosomal proteins were extracted from the open reading frames (ORFs) in the CPR genomes, and 443– 890 sequences for each ribosomal protein were obtained. Ribosomal protein sequences of non-CPR bacteria were extracted according to the annotation in RefSeq, and 1,454–2,613 sequences were obtained for each ribosomal protein.

Figure 2 shows the presence or absence of each ribosomal protein in the 69 complete genomes of CPR bacteria and 140 representative non-CPR bacteria (70 free-living lineages and 70 symbiotic lineages). Consistent with a previous report (Brown et al. 2015), no L30 was detected in the CPR bacteria. L30 is reportedly encoded in a gene cluster and is located between S5 and L15 in many bacteria (Cerretti et al. 1983; Roberts et al. 2008), but S5 and L15 are located close to each other in CPR bacterial genomes (data not shown). We also confirmed that L1 is absent in a subgroup of Parcubacteria, and that L9 is absent in Microgenomate (excluding Beckwithbacteria), Saccharibacteria (phylum group I), and WWE3 and Dojkabacteria (phylum group II).

**Figure 2.**
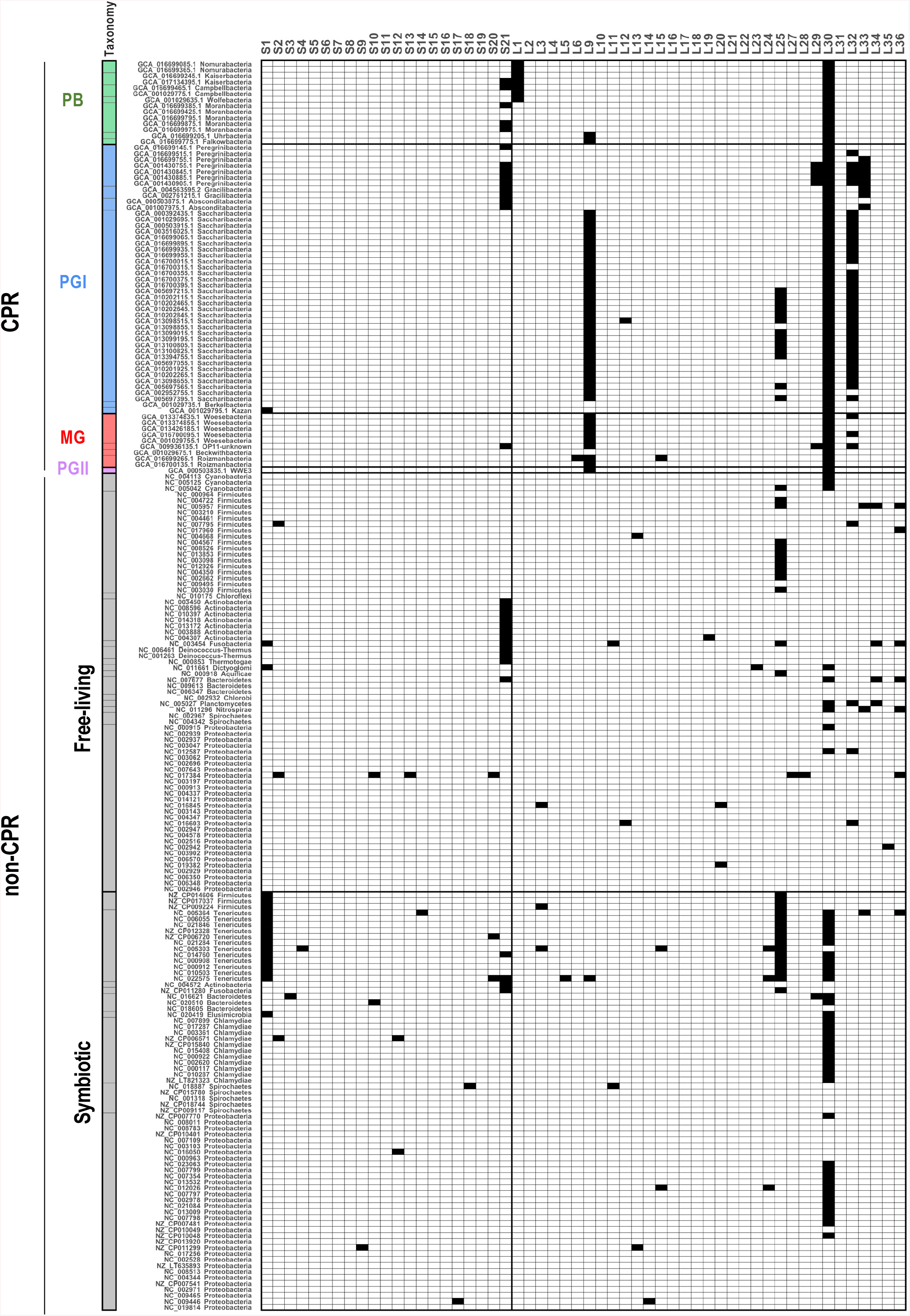
Presence and absence of ribosomal proteins encoded in complete CPR and non-CPR genomes. The distributions of 54 ribosomal proteins (columns) across representative bacterial genomes (rows) are shown. Presence (white) or absence (black) of each ribosomal protein is indicated for 69 complete genomes of CPR bacteria and 140 genomes of representative non-CPR bacteria (70 free-living lineages and 70 symbiotic lineages) (see Supplementary Tables S2–S4). The rows were sorted based on a phylogenetic tree (Hug et al. 2016), and the labels provide NCBI accession numbers and the taxonomy of each genome. Left panel shows the phylum-level classification and is colored according to taxonomic group (see Supplementary Figure S1): Parcubacteria (PB): green; Microgenomates (MG), red; phylum group I (PGI), blue; phylum group II (PGII), purple; non-CPR, light gray (see Supplementary Figure S1).

New ribosomal protein deficiencies were also found in recently reported complete CPR genomes. Clades containing Peregrinibacteria, Gracilibacteria, and Absconditabacteria, belonging to phylum group I, frequently lacked S21 and L33, and some Peregrinibacteria also lacked L29 and L32. Saccharibacteria usually lacked L32, except for one complete genome. About half the phyla also lacked L25. Although the use of complete genomes is suitable for assessing the loss of specific genes encoding ribosomal proteins, the number of CPR bacterial lineages for which complete genomes are available is currently limited, and 65% of the complete genomes used in the present study are concentrated in phylum group I (Supplementary Table S1A). Therefore, to investigate the distribution of ribosomal proteins throughout the CPR bacteria, the frequency of loss of each ribosomal protein in each phylum was also calculated using all the data, including both the complete genomes and the draft genomes (Supplementary Figure S3). Here, we targeted all CPR phyla for which three or more genomes could be obtained (either complete or draft). The ribosomal proteins which were not detected in more than 80% of the genomes from any of the phyla were then extracted. The results showed that S21 was also frequently missing in Pacubacteria, and Collierbacteria of Microgenomates, as in ‘phylum group I’ (as mentioned above). L28 was lacking in all draft genomes of Daviesbacteria of Microgenomates.

In contrast, most ribosomal proteins were widely conserved in the non-CPR bacterial genomes, although ribosomal proteins S1, S21, L25, and L30 were missing across at least two non-CPR phyla (Figure 2 and Supplementary Figure S3). These ribosomal proteins have already been reported as readily lost in non-CPR bacteria (Yutin et al. 2012; Grosjean et al. 2014). Mollicutes, a phylum of symbiotic bacteria, is particularly deficient, and always lacked at least two (and up to eight) ribosomal proteins.

In summary, S21, L25, and L30 were frequently missing in both CPR and non-CPR, and moreover L1, L9, L28, L29, L32, and L33 were also missing preferentially in CPR. It seems that LSU proteins are more likely to be deficient in CPR than are SSU proteins. Although these proteins are widely distributed in non-CPR, any ribosomal protein can be missing in specific non-CPR bacterial genomes, particularly in small genomes (Lecompte et al. 2002; Nikolaeva et al. 2021).

### Many ribosomal proteins in CPR bacteria differ from those in non-CPR bacteria

We analyzed the presence or absence of ribosomal proteins in CPR bacteria and clarified the lineage-specificity of missing ribosomal proteins. Because the molecules that make up the ribosome interact in a complex manner, it is possible that the lack of individual specific ribosomal proteins could cause changes in these other proteins. Therefore, we examined the size and amino acid sequences of individual ribosomal proteins in CPR bacteria and compared them with those in non-CPR bacteria. Figure 3 shows the size distributions and amino acid sequence alignments of three ribosomal proteins, L13, S19, and L1, as examples of significantly different sequences in CPR and non-CPR bacteria. Although the L13, S19, and L1 proteins each show constant size distributions in most species of non-CPR bacteria, the size distributions of these proteins in CPR bacteria have two peaks, and the proteins in one group are significantly smaller (L13) or larger (S19 and L1) than each corresponding protein in non-CPR bacteria (Figures 3A–3C). When these amino acid sequences were aligned, regions missing in particular lineages of CPR bacteria and regions present in only some CPR bacteria were detected. Because genes encoding ribosomal protein genes of abnormal length are often found in CPR bacterial genomes as a result of sequencing errors or misassembly during the cloning and gene identification steps, we focused only on features that are shared among closely related species. The ribosomal protein L13 tends to lack the N-terminal (approximately 12 aa) and C-terminal (approximately 16 aa) regions in most of the Parcubacteria (Figure 3D). In the C-terminal region of the ribosomal protein S19, an alanine- and lysine-rich region (approximately 25 aa) is specifically present in Parcubacteria and phylum group I (Figure 3E). As mentioned above, the ribosomal protein L1 is absent in a group of Parcubacteria (Brown et al. 2015) (Figure 2 and Supplementary Figure S3), and the aa sequences of L1 proteins in lineages other than Parcubacteria also have characteristics that differ from those in non-CPR bacteria (Figure 3F). In Berkelbacteria (phylum group I) and in Microgenomates other than Woykebacteria, a few internal regions of the L1 protein (approximately 20–40 aa, in total) are absent, and a specific region of about 40–130 aa is inserted at the N-terminus. This N-terminal region is also present in Saccharibacteria (phylum group I) and Woykebacteria, and some members of WWE3 (phylum group II), in which the internal region is not missing. The C-terminal region of the L13 protein and the internal region of the L1 protein that are missing in CPR bacteria correspond to regions containing highly conserved aa residues in non-CPR bacteria. In contrast, the CPR-specific regions inserted at the C-terminus of the S19 protein and the N-terminus of the L1 protein do not match any known functional domain registered in the Pfam database (Mistry et al. 2021).

**Figure 3.**
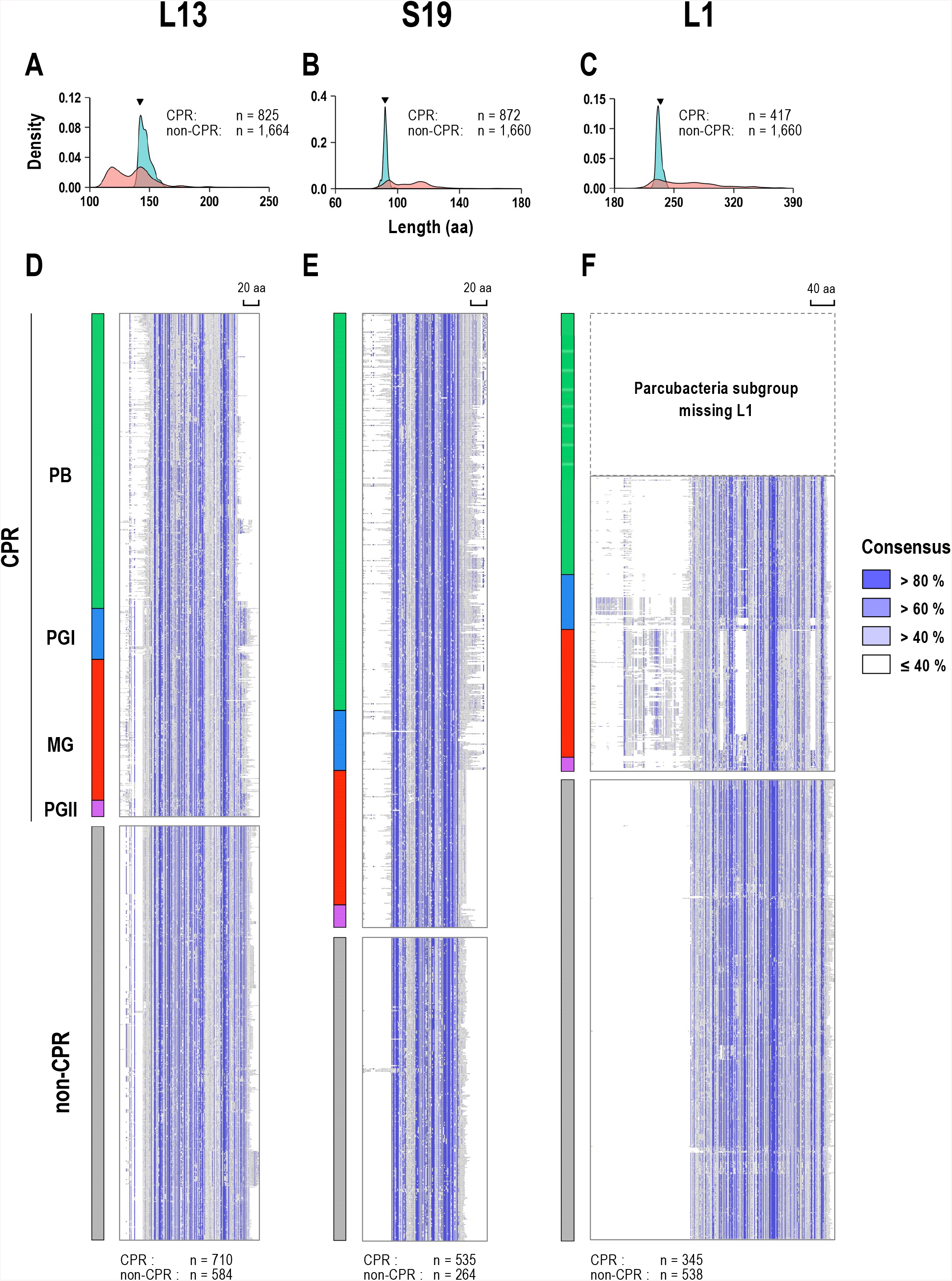
Multiple amino acid sequence alignments of three selected ribosomal proteins that differ in length in CPR and non-CPR bacteria. (A–C) Distributions of the lengths of three ribosomal proteins (A: L13; B: S19; and C: L1) with distinctly different lengths in CPR and non-CPR bacteria. Density curves for each protein length were calculated based on all the sequences in the dataset (CPR: light red; non-CPR: light blue). The total number of sequences is indicated in each panel. The length of each ribosomal protein in *E. coli* is indicated with a black inverted triangle. (D–F) Multiple amino-acid-sequence alignments of three ribosomal proteins corresponding to panels A–C (D: L13; E: S19; and F: L1) are shown in the order based on a phylogenetic tree (Hug et al. 2016). Representative sequences were selected from each CPR and non-CPR group, and poorly aligned columns were removed (see Materials and Methods). The consensus residues in each column are highlighted in blue gradient, according to the percentage identity calculated when gaps were ignored. The magnification and aspect ratio of each alignment panel have been adjusted for clarity. The scale bars of the alignment columns (aa lengths) are shown at top right. The number of sequences included in the alignment is shown at the bottom. Panels on the left side of the alignment are colored according to the taxonomic group of each sequence (see Supplementary Figure S1): Parcubacteria (PB) in green, Microgenomates (MG) in red, phylum group I (PGI) in blue, phylum group II (PGII) in purple, and non-CPR in light gray.

Supplementary Figure S4 shows the length distributions and aa sequence alignments of 53 ribosomal proteins, excluding the L30 protein, which is completely absent from CPR bacterial genomes. Surprisingly, in about half the ribosomal proteins, as well as the abovementioned three ribosomal proteins, the positions and numbers of peaks in the size distributions differ significantly between CPR and non-CPR bacteria. Ribosomal proteins S1, S3, S14, L2, L3, L5, L15, and L20 tend to be smaller in CPR bacteria. In particular, similar to the L13 protein (Figures 3A and 3D), L2, L3, L5, and L15 proteins are significantly smaller in some CPR bacteria than in non-CPR bacteria, and these small proteins lack an N-terminal (L2 and L5 proteins), C-terminal (L15 protein), or internal region (L3 protein). The S1, S3, S14, and L20 proteins have various patterns of length in non-CPR bacteria, but in most CPR bacteria they are similar in size to the smallest group of proteins in non-CPR bacteria. Among these proteins, S1 is known to consist of a different number of S1 domains, depending on the species (Machulin et al. 2019). In non-CPR bacteria, S1 proteins with six domains account for about 60% of the total known S1 proteins, whereas in many CPR bacteria, most S1 proteins are composed of 3–4 S1 domains (Supplementary Figure S4). The ribosomal proteins with regions that are specifically missing in CPR bacteria were L1, L2, L3, L5, L13, and L15. These proteins occur preferentially in the LSU, which is also true of the completely absent proteins in CPR bacteria (Figure 2). In addition to the S19 (Figure 3B and 3E) and L1 proteins mentioned above (Figure 3C and 3F), many ribosomal proteins are large in the CPR bacteria: S6, S10, S11, S12, S13, S15, S21, L12, L19, L22, L23, L25, L27, L31, and L32. In particular, regions specifically present in only a proportion of CPR bacteria were detected in the C-terminal regions of the S13 and S15 proteins, in the N-terminal region of the L23 protein, and in the internal region of the L27 protein. The internal region of S12, which only occurs in Tenericutes, Chloroflexus, and some Firmicutes in non-CPR bacteria, is present in almost all S12 sequences of CPR bacteria. Similar to the L1 protein (Figure 3C and 3F), the S20 protein aa sequence has both a missing region (center) and a specific extra (C-terminal) region. However, in the case of the proteins with additional N-terminal region, it should be noted that the start codon of the ORF was predicted based on the location of the first start codon, and is not necessarily the actual start codon. Although the folding structures and functions of the CPR-specific protein regions have not been characterized, in some examples the C-terminal extension of the ribosomal proteins appears to improve the stability of rRNA folding, and also contributes to the environmental adaptation of *Thermus thermophilus* (Melnikov et al. 2018).

### Smaller rRNAs and intron-containing rRNAs in CPR bacteria

Ribosomal RNAs form the basis of the ribosome and play a major role in translation (Nissen et al. 2000). By comparing the sizes of the rRNA genes in CPR bacteria and non-CPR bacteria, we found that the rRNAs in CPR bacteria are rather small. With an Infernal search using rRNA secondary structure models, 375 full-length genes for 16S rRNA, 347 full-length genes for 23S rRNA, and 630 full-length genes for 5S rRNA were obtained from the complete and draft genomes of CPR bacteria. The CPR bacteria basically had one copy of each rRNA gene per genome. Figure 4A–4D shows the distribution of rRNA gene lengths in CPR bacteria, symbiotic non-CPR, and free-living non-CPR bacteria. As reported previously (Brown et al. 2015), some CPR bacteria have long 16S and 23S rRNA genes, containing insertion sequence(s) (Figure 4A and 4B). In particular, we found that most 23S rRNAs in CPR bacteria contain insertion sequences, ranging from 0.5 kb to several kilobases in total length. When the insertion sequences were extracted, based on alignments with the rRNA genes of *E. coli* K-12, 42% of the 16S rRNA genes and 77% of the 23S rRNA genes of the CPR bacteria contained long insertions of ≥100 bases. The average length of insertions per gene was 623 bases (maximum 5.5 kb) for the 16S rRNA gene and 1,232 bases (maximum 5.7 kb) for the 23S rRNA gene. Because the inserted regions were estimated by comparison with *E. coli* rRNAs, they do not exactly reflect the regions excluded after RNA splicing, but it is obvious that the 16S and 23S rRNA genes of CPR bacteria frequently contain insertions. The length profile of the 5S rRNA in CPR bacteria closely resembles that of parasitic non-CPR bacteria, with two peaks at 105 and 120 bases, whereas the peak in the 5S rRNA of free-living non-CPR bacteria is predominantly at 120 bases (Figure 4C). Therefore, many of the 5S rRNAs in CPR bacteria are much smaller than those in free-living non-CPR bacteria.

**Figure 4.**
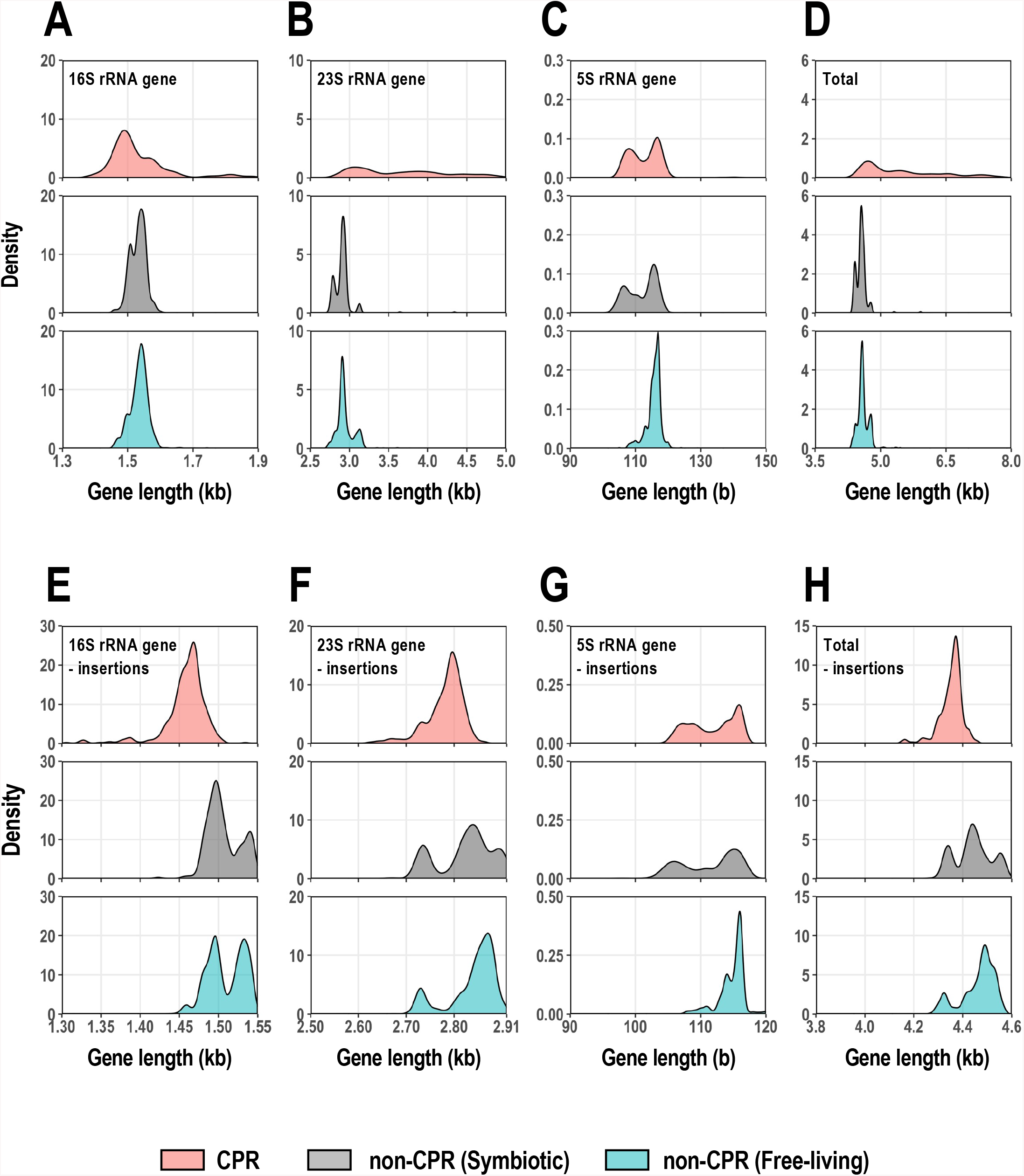
Length distributions of rRNA genes encoded in CPR and non-CPR genomes. (A–D) Density distributions of the whole-length 16S, 23S, and 5S rRNA genes and the total combined lengths of these three genes. Each horizontal axis is limited to the range in which the main peak occurs, and a few genes beyond this limit (A: 5.7%; B: 3.0%; C: 0.3%; and D: 1.5% of the total data) were omitted from the figures. (E–H) Density distributions of 5S, 16S, and 23S rRNA gene length excluding insertion sequences, and their combined length for each genome. The numbers of data were: 16S rRNA genes (CPR bacteria, n = 375; symbiotic non-CPR bacteria, n = 167; free-living non-CPR bacteria, n = 1,494); 23S rRNA genes (CPR bacteria, n = 347; symbiotic non-CPR bacteria, n = 167; free-living non-CPR bacteria, n = 1,494); and 5S rRNA genes (CPR bacteria, n = 630; symbiotic non-CPR bacteria, n = 167; free-living non-CPR bacteria, n = 1,494) and the three genes combined (CPR bacteria, n = 240; symbiotic non-CPR bacteria, n = 167; free-living non-CPR bacteria, n = 1,494). Density curves are colored according to the group (CPR, light red; symbiotic non-CPR, light gray; free-living non-CPR, light blue).

We removed the insertion sequences from each CPR rRNA gene based on a comparison with the corresponding *E. coli* rRNA genes to roughly estimate the length distributions of the mature rRNAs (after RNA processing) in CPR bacteria (Figure 4E–4G). The distributions of rRNA lengths in non-CPR bacteria had 2–3 peaks for all rRNA types, and the proportion of shorter RNAs was greater in symbiotic bacteria than in free-living bacteria. In contrast, the size distributions of the 16S and 23S rRNA genes (without insertion sequences) in the CPR bacteria mainly had one large peak. The standard size (approximately 1.47 kb) of 16S rRNA in CPR bacteria, estimated from the location of the peak, was about 30 bases smaller than size of smaller 16S rRNA gene group in non-CPR bacteria (Figure 4E). However, most of the 23S rRNAs in CPR bacteria had an intermediate size (approximately 2.80 kb), between the two peaks found in the non-CPR bacteria. This was about 50 bases smaller than the larger 23S genes in the non-CPR bacteria (peak on the right in Figure 4F), which accounted for the majority of non-CPR bacteria, but non-CPR also included smaller 23S rRNA genes, at around 2.72 kb (peak on the left in Figure 4F). Among the non-CPR bacteria, these very small 23S rRNAs were abundant in the Proteobacteria, whereas among the CPR bacteria, Magasanikbacteria of Parcubacteria also had small 23S rRNAs. The size distribution of the 5S rRNAs in CPR bacteria displayed two peaks, similar to the distribution of parasitic bacteria, and the smaller gene group, which accounted for about half the total genomes, had 105–110 bases and mainly occurred in Parcubacteria (Figure 4G). In free-living non-CPR bacteria, the proportion of 5S rRNA genes with ≤110 bases was only 8%. Summing the lengths of the three rRNA genes and excluding each insertion sequence, the most frequent lengths were in the following order: CPR < symbiotic non-CPR < free-living non-CPR (*p* < 0.01 for each pair, Bonferroni’s test) (Figure 4H). The most frequent value for the total rRNA length in CPR bacteria (4.37 kb) was 118 bases shorter than that in the non-CPR bacteria. The total rRNA length of some non-CPR bacteria was the same as that of CPR bacteria, but more often in symbiotic bacteria than in free-living bacteria. These results suggest that the core region of rRNA is smaller in CPR bacteria than in typical non-CPR bacteria.

To investigate the smaller rRNA genes of the CPR bacteria at the nucleotide sequence level, a multiple alignment analysis of CPR and non-CPR rRNA genes was performed. Using the *E. coli* rRNA genes as the reference sequences, the insertion sequences were removed from each rRNA gene of the CPR bacteria, and the resulting nucleotide sequences of the CPR and non-CPR rRNA genes were compared. The results showed that all three types of rRNA genes in the CPR bacteria frequently lacked one region (in 5S rRNA) or several regions (in 16S and 23S rRNAs) (Supplementary Figure S5). The 16S rRNA gene of CPR bacteria has five gap regions (the regions with higher gap rates in CPR bacteria than in non-CPR bacteria were designated 16Gap1–16Gap5), and the number of non-CPR bacterial lineages lacking regions 16Gap4 and the 16Gap5 was extremely limited. The 16Gap4 region was deleted in the phylum Chloroflexi and in part of the class Tenericutes, which contains symbiotic bacteria, and the 16Gap5 region was deleted in the phyla Bacteroidetes and Chlorobi. Additionally, although it has been reported that the anti-SD sequence at the 3’ end of the 16S rRNA is rarely lost (Li et al. 2012; Amin et al. 2018), in our data, the anti-SD motif (CCTCCT) (Nikolaeva et al. 2021) was detected less frequently in some lineages of Parcubacteria (detection rate in Parcubacteria of 47.8%). The lineages without anti-SD motif overlapped with the many of L1-missing group (Figure 2 and Supplementary Figure S3), but the functional association between the anti-SD sequence and L1 is unclear. In the 23S rRNA genes, four regions (designated 23Gap1–23Gap4) are deleted throughout the CPR bacteria or in a lineage-specific manner. The 23Gap3 region is deleted in some Parcubacteria (e.g., Magasanikbacteria, Uhbacteria, and Falkowbacteria), Microgenomates and phylum group II in CPR bacteria, but only in two species of non-CPR bacteria. The 5S rRNA genes of CPR bacteria have one gap region (designated 5Gap1), which is deleted in many Parcubacteria and some non-CPR bacteria. Unexpectedly, while there are these missing regions in rRNA genes of CPR bacteria, a 16S rRNA gene region corresponding to nucleotides 837–850 of the E. coli gene was highly conserved in CPR bacteria, especially in Parcubacteria and phylum group I, although it was weakly conserved in the non-CPR bacterial dataset and is reported to be commonly lost in small genomes (Nikolaeva et al. 2021).

Mapping the missing regions in each rRNA gene of the CPR bacteria against the known *E. coli* rRNA secondary structures revealed that all gap regions correspond to the entire or the tip of particular stem–loop structures in each type of rRNA (Figure 5A). For example, the lack of 16Gap5 in the CPR bacteria indicates that the tip of helix 44 is lost and the helix slightly shortened. Although helix 44 contributes to the accuracy of translation initiation (Qin et al. 2012), the position of 16Gap5 does not affect known functional sites. 5Gap1 in the 5S rRNA corresponds to helix IV and loop D, a region that shows large structural variation in bacteria, and can be lost (Szymanski et al. 2016; Stepanov and Fox 2021). Some rRNA helices are known to be deleted in small non-CPR genomes (Nikolaeva et al. 2021), but no case of 23SGap3 (the loss of 23S rRNA helix 78) has yet been reported. Therefore, the rRNAs of CPR bacteria are seen to have structures in which multiple stem–loops are lacking or shortened relative to the corresponding *E. coli* rRNA structures. In contrast, although all CPR bacteria lack L30, they retain the sequence encoding the region in the vicinity of the loop E, which contains the L30 binding site for 5S rRNA (Sun and Caetano-Anolles 2009). The 16S rRNA region highly conserved in Parcubacteria and phylum group I, (surrounded by dashed lines in Figure 5A) was corresponds to helix h26, which interacts with the SD helix (base pairs with the SD sequence in mRNAs and the anti-SD sequence of 16S rRNA) and is considered to be contributes to the start of translation (Korostelev et al. 2007). It is thought that the SD helices do not form in some Parcubacteria because they lack the anti-SD motif, but the role of conserved helix h26 is unknown.

**Figure 5.**
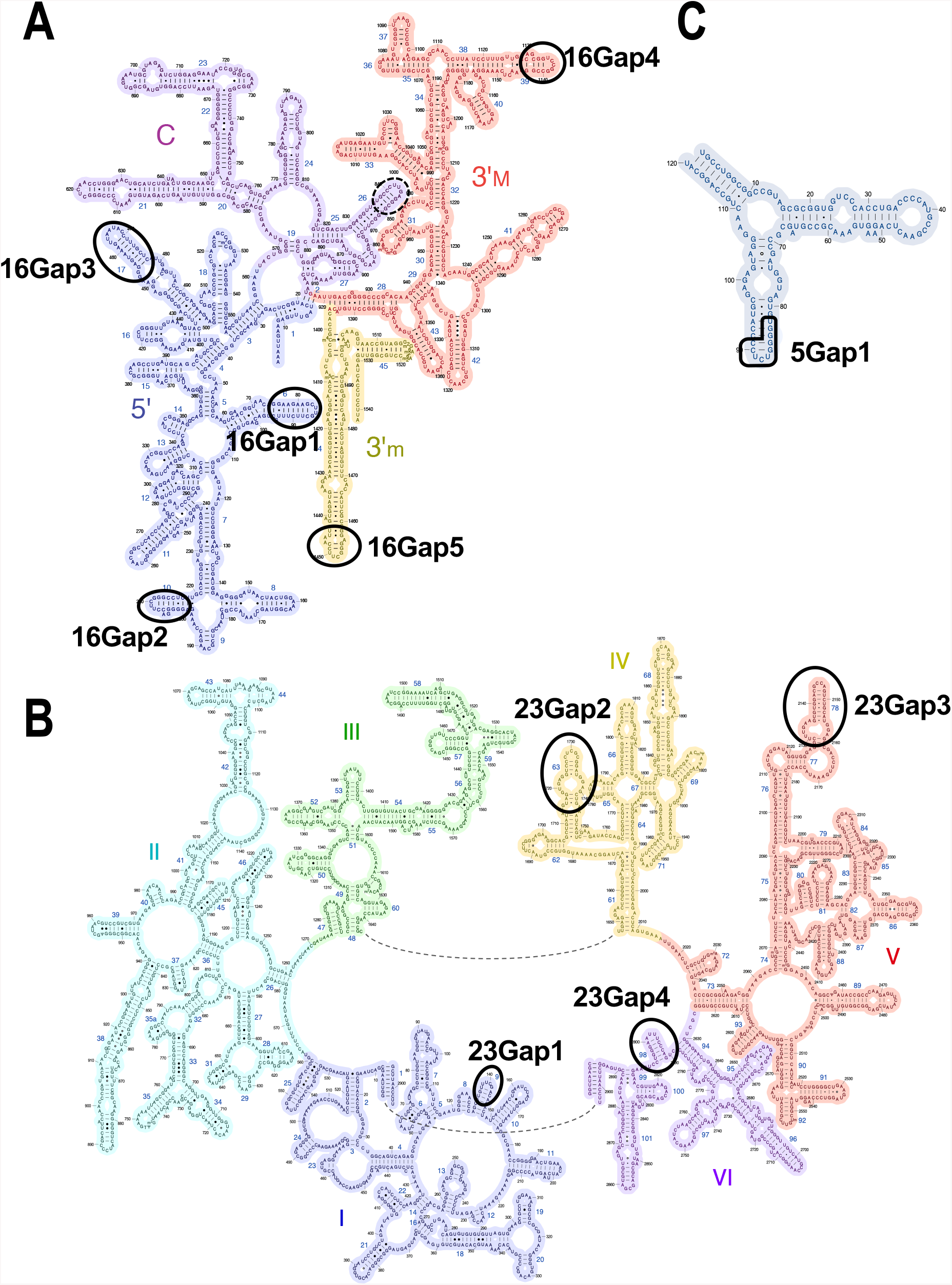
rRNAs of CPR bacteria lack terminal stem–loop domain(s) in their RNA secondary structures. RNA regions lacking in CPR bacteria are indicated by ellipses in the secondary structures of (A) 16S, (B) 23S, and (C) 5S rRNAs of *E. coli*. The dashed line indicates a region highly conserved in CPR bacteria. See Supplementary Figure S7 for details. These RNA secondary structures were obtained from the Center for Molecular Biology of RNA (http://rna.ucsc.edu/rnacenter/ribosome_images.html).

### Ribosomes of CPR bacteria lack RNA and protein regions present on the ribosomal surface

To roughly estimate the shape of CPR bacterial ribosomes, the ribosomal proteins and rRNA regions missing in all or some CPR bacteria were mapped onto a well-studied ribosomal structural model of *E. coli* strain K-12 (Figure 6A and 6B). Ribosomal proteins bind around the rRNA backbone to form the outer part of the ribosome. As mentioned above, most of the ribosomal proteins lacking in the CPR bacteria occur in the large subunit, and all but the L33 protein are exposed on the ribosomal surface, and the L1, L9, and L28 proteins are located close to each other on the ribosome surface (Figure 6A). Although rRNAs form the core of the ribosome, the regions in the 16S and 23S rRNAs that are lacking in CPR bacteria (Supplementary Figure S5) are all exposed on the surface of the ribosome (Figure 6B). Moreover, the absent regions in the 16S rRNA (16Gap1, -2, and -3, corresponding to helices h6, h10, and h17, respectively) and the absent regions in the 23S rRNA (23Gap1 and -2, corresponding to helices h9 and h63, respectively) are located close to each other in the ribosome tertiary structure, suggesting that the local structures formed by these helices are lost on the surface of the ribosome in CPR bacteria. When we compared the missing parts of the rRNAs and ribosomal proteins, L1 and 23Gap3 (helix 78), L32 and 23Gap4 (helix 98), and L25 and 5Gap1 were located close to each other, respectively. (Figure 6C). The L1 protein and 23S rRNA helix 78 are known to form a mobile structure called the ‘L1-stalk’, which contributes to translation efficiency (Trabuco et al. 2010; Reblova et al. 2012), but both are specifically absent in some CPR bacteria (Supplementary Figure S5). Although L1 and 23Gap3 are not always lacking in the same lineages, in at least some CPR bacterial ribosomes the L1-stalk is absent or incomplete. Some ribosomal proteins have regions that are specifically present in only some CPR bacterial lineages (Figure 3 and Supplementary Figure S4). However, it is unclear whether these regions compensate for the missing regions found in these bacteria.

**Figure 6.**
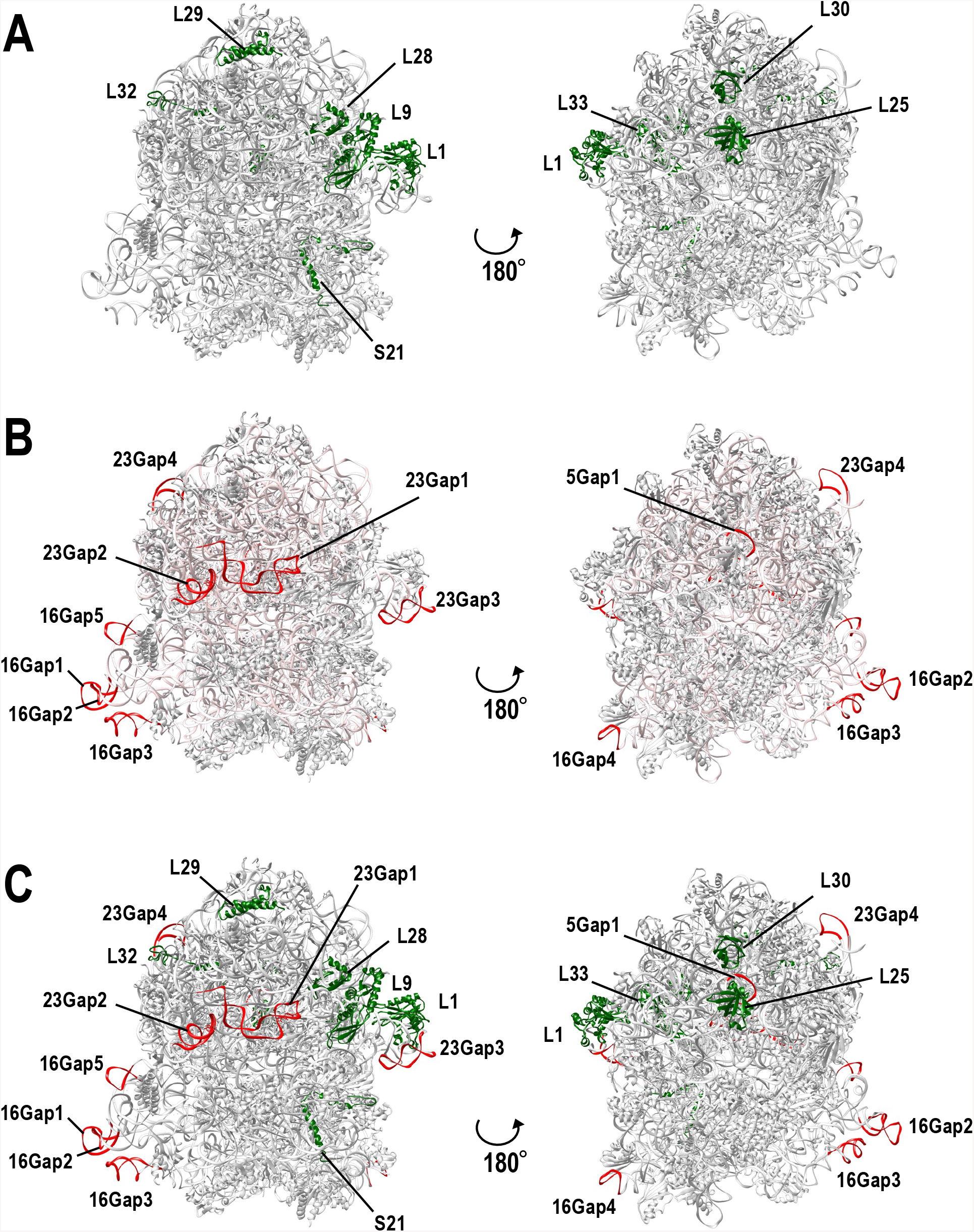
Missing rRNA regions and missing ribosomal proteins in CPR bacteria map to the surface of the 3D *E coli* ribosomal structure. A three-dimensional structural model of *E. coli* K-12 (PDB ID: 5U9G) ribosome was used for the analysis. (A) Ribosomal proteins missing in some or all CPR bacteria (see Figure 2 and Supplementary Figure S3) are shown in green. (B) 16S, 23S, and 5S rRNA regions missing in CPR bacteria (see Figure 5) are colored red. Entire rRNA regions are colored light pink. (C) Mixed view of (A) and (B).

In this study, we detected the lack of several ribosomal regions in all or some lineages of CPR bacteria. These missing regions are biased to the outside surface of the ribosomal complex and are thought to simplify the surface structure of the ribosomes of CPR bacteria. It has been reported that the evolution of ribosomes (i.e., the expansion of rRNA molecules and the acquisition of new ribosomal proteins) has progressed from the center of the ribosome to the outer surface. Previous studies have proposed that the evolution of the ribosome progressed in six phases, based on predictions of the order of acquisition of each segment of the prokaryotic rRNA (Petrov et al. 2014; Petrov et al. 2015). According to their theory, the rRNAs in the SSU and LSU evolved independently from phases 1 to 3, and that interaction between the subunits was formed in phases 3 and 4. In phase 5, the acquisition of functional ribosomal proteins began with the integration of the 5S rRNA. The ribosomal proteins strengthened the binding between the subunits and formed the binding sites for translation factors. In the final phase 6, the rRNA regions located on the ribosome surface were acquired, and the surface was covered with proteins that bound to the regions of the rRNAs (proteinizing). In this evolutionary model, all the regions missing in the CPR rRNAs (Figure 5 and Supplementary Figure S5) are considered to have been acquired in phase 5 or 6. The 16S rRNA region (nucleotides 837–850 in *E. coli* 16S rRNA) that shows much greater conservation in CPR than in non-CPR was also acquired in phase 6. In contrast, the detailed order of acquisition of the ribosomal proteins has not yet been predicted, but except for L33, the ribosomal proteins that are lacking in CPR are exposed on the ribosome surface when they are present in other bacteria (Figure 6A), so it is thought that many of them were acquired in phase 6. The central regions of the ribosome, which formed in the early stage of molecular evolution, are conserved throughout the three domains of life (Bacteria, Archaea, and Eukaryota). They play a central role in translation (Melnikov et al. 2012; Bernier et al. 2018), whereas the surface regions acquired in the late stage contribute to the efficiency of translation and the stability of the ribosome, and are not essential in many cases (Galperin et al. 2021a). Because the central regions of the ribosome are strongly conserved, even in CPR bacteria, the lack of certain molecules (described in this paper) is thought to simplify the structure of the ribosome because their absence does not basically affect the regions essential for translation. Our study provides concrete examples that support the theory of ribosome evolution.

## MATERIALS AND METHODS

### Data sources

We downloaded 897 publicly available genomes of CPR bacteria (69 complete and 828 draft genomes) from GenBank at the NCBI site (https://ftp.ncbi.nlm.nih.gov/genomes/genbank/) (Sayers et al. 2019) (Supplementary Tables S1A and S2). These genomes were classified into four subgroups based on a previously proposed phylogenetic tree (Hug et al. 2016): Parcubacteria and Microgenomates, which are proposed superphyla, and phylum group I and phylum group II, which are temporarily designated groups in this study (Supplementary Figure S1). As the control, we also downloaded 1,661 complete genomes of non-CPR bacteria, described as “Reference” or “Representative” in the NCBI Reference Sequence Database (RefSeq) (https://ftp.ncbi.nlm.nih.gov/genomes/refseq/; accessed on November 1, 2018; (O’Leary et al. 2016). These non-CPR bacterial genomes were classified into endosymbiotic or parasitic groups (symbiotic non-CPR bacteria, n = 167) and others (free-living non-CPR bacteria, n = 1,494) (Supplementary Table S1B). Seventy genomes from each of these two non-CPR groups were selected as representative (Supplementary Tables S3–S4). Protein-coding genes in the CPR genomes were predicted with Prodigal v2.6.3 (Hyatt et al. 2010) to analyze the protein length distributions (Figure 1 & Supplementary Figure S2). The genes of Absconditabacteria and Gracilibacteria were predicted with genetic code 25 (Campbell et al. 2013), in which the UGA stop codon is translated as glycine, and the genes of the other genomes were predicted with the standard bacterial genetic code.

### Estimation of genome sizes in CPR bacteria

The genome sizes of the CPR draft genomes were estimated by assessing their completeness using 43 universal single-copy genes (SCGs) for CPR bacteria (Brown et al. 2015). Each CPR genome was translated in six frames with the getORF program in EMBOSS 6.6.0 (Rice et al. 2000). Genetic code 25 was used for Absconditabacteria and Gracilibacteria (Campbell et al. 2013), and code 11 was used for the other taxa. All possible ORFs with a minimum length of 10 amino acids were extracted for subsequent analysis. Using SCG proteins from the NCBI Clusters of Orthologous Groups of proteins (COG) database (https://www.ncbi.nlm.nih.gov/COG/) (Galperin et al. 2021b) as queries, BLASTP searches (blast+ version 2.9.0) (Camacho et al. 2009) were performed against the ORFs of CPR bacteria (*E*-value threshold: 1*e*−5) to identify SCG candidates. These candidate SCG proteins were then subjected to a reverse BLASTP search against the same SCG protein set from the COG database, and the top hits were used to confirm the assignments. The ratio of SCGs detected in each genome was defined as the genome restoration rate, and the estimated genome size was obtained by dividing the total length of the scaffold by the genome restoration rate.

### Search for ribosomal protein genes and rRNA genes

The ribosomal protein genes and rRNA genes in CPR bacterial genomes were detected with sequence similarity searches. Ribosomal protein sequences were compared with the ORFs in the CPR bacterial genomes (see the previous section for detecting ORFs), using a combination of BLASTP and hmmscan. Using the amino acid sequence sets of 54 bacterial ribosomal proteins from the NCBI COG database as the queries (Supplementary Table S5), BLASTP searches (blast+ version 2.9.0) (Camacho et al. 2009) were performed against the ORFs in the CPR bacterial genomes (*E*-value threshold: 1*e*™5; query cover threshold: 50%) to identify candidate ribosomal proteins. These candidate proteins were then subjected to a reverse BLASTP search against the same set of 54 bacterial ribosomal proteins from the COG database. The top hits obtained confirmed the assignments. The ORFs from the CPR bacterial genomes were then compared with the Pfam ribosomal protein HMM profiles (Mistry et al. 2021), using hmmscan from the HMMER 3.3.1 package (Eddy 2011). A set of ribosomal protein sequences was generated by combining the sequences found with these two methods, and was inspected to remove false positive hits, particularly those observed in targets containing ubiquitous RNA-binding domains.

To detect 16S, 23S, and 5S rRNA sequences in the CPR genomes, the cmsearch program from the Infernal package (version 1.1.3) was used (*E*-value threshold: 1*e*−4) (Nawrocki and Eddy 2013). Here, we used RNA secondary structure models for 5S rRNA (RF00001), 16S rRNA (RF00177), and 23S rRNA (RF02541) obtained from the Rfam database (http://rfam.xfam.org/) (Kalvari et al. 2018). Because CPR bacteria often have long insertions within their 16S and 23S rRNA genes, several partial hits were identified. If the partial hits were adjacent on the same scaffold (i.e., the gaps between hits were ≤5,000 bases and no hits for other rRNAs were identified in the gap), those hits were considered to be single genes. Ribosomal protein and rRNA genes in the non-CPR bacteria were identified according to the RefSeq annotation, and one representative sequence per genome was extracted for each protein and rRNA. Because one non-CPR genome (*Streptococcus pyogenes*, accession: NC_002737) lacked annotation of the 5S rRNA gene, it was evaluated with cmsearch. The partial gene sequences truncated at the end of the scaffold were removed from the subsequent analysis. The sequences of the CPR and non-CPR rRNA genes obtained were aligned with MAFFT L-INS-i (v7.407) (Katoh and Standley 2013), and the insertion sequences were identified based on a comparison with the well-studied *E. coli* rRNA genes (16S, 1,542 bases; 23S, 2,904 bases; 5S, 120 bases), which were included in the non-CPR dataset.

### Comparative sequence analysis and structural mapping

Representative sequences of ribosomal proteins and rRNA genes were selected for multiple alignment and visualization. After sequences containing unknown amino acids (‘X’) or unknown nucleotides (‘N’) were removed, the genes were clustered based on sequence identity using the UCLUST algorithm (cluster_fast command) in USEARCH v11 (Edgar 2010), and the cluster centroids (typical sequences) were selected as representative. The identity thresholds were 80% for the ribosomal proteins, 5S rRNA genes, and 23S rRNA genes, and 85% for the 16S rRNA genes, based on the number of clusters generated. The representative sequences were aligned with MAFFT L-INS-i (v7.407) (Katoh and Standley 2013). To visualize the alignment of ribosomal protein sequences, columns with gap frequency of >90% in both of the CPR and non-CPR groups were removed. To visualize the rRNA gene sequence alignment, insertions with respect to the *E. coli* K-12 genes were removed. The alignments were visualized with Jalview 2.11 (Waterhouse et al. 2009). The locations of the missing regions in the CPR bacterial ribosomal proteins and rRNAs were estimated by mapping the genes and rRNAs onto the tertiary structure of the *E. coli* K-12 ribosome (PDB ID: 5U9G) (Demo et al. 2017). Three-dimensional mapping was performed with UCSF Chimera (Pettersen et al. 2004).

## ACKNOWLEDGMENTS

The authors thank Dr. Shigenori Maruyama for his critical suggestions. We also thank all the members of the RNA Group at the Institute for Advanced Biosciences of Keio University, Japan, for their insightful discussions. This work was supported, in part, by a KAKENHI Grant-in-Aid for JSPS Fellows (21J12231) and research funds from the Yamagata Prefectural Government and Tsuruoka City, Japan. The funding bodies played no role in the study design, data collection or analysis, the decision to publish, or the preparation of the manuscript.

## CONFLICT OF INTEREST STATEMENT

The authors declare that they have no conflicts of interest.

## SUPPLEMENTARY INFORMATION

Supplementary material is available for this article: Supplementary Tables S1–S6 and Supplementary Figures S1–S5.

## Notes

### Competing Interest Statement

The authors have declared no competing interest.

